# Gait-phase modulates alpha and beta oscillations in the pedunculopontine nucleus

**DOI:** 10.1101/2021.03.05.434086

**Authors:** Shenghong He, Alceste Deli, Petra Fischer, Christoph Wiest, Yongzhi Huang, Sean Martin, Saed Khawaldeh, Tipu Z. Aziz, Alexander L Green, Peter Brown, Huiling Tan

**Affiliations:** MRC Brain Network Dynamics Unit at the University of Oxford, United Kingdom; Nuffield Department of Clinical Neurosciences, University of Oxford, United Kingdom.; Nuffield Department of Surgical Sciences, University of Oxford, United Kingdom.; Academy of Medical Engineering and Translational Medicine, Tianjin University, China; Oxford Centre for Human Brain Activity, Wellcome Centre for Integrative Neuroimaging, University of Oxford, United Kingdom

**Keywords:** Pedunculopontine nucleus (PPN), Parkinson’s disease, Multiple System Atrophy, freezing of gait, deep brain stimulation, gait-phase related modulation

## Abstract

**Backgroud:** The pedunculopontine nucleus (PPN) is a reticular collection of neurons at the junction of the midbrain and pons, playing an important role in modulating posture and locomotion. Deep brain stimulation of the PPN has been proposed as an emerging treatment for patients with Parkinson’s disease (PD) or multiple system atrophy (MSA) suffering gait-related atypical parkinsonian syndromes.

**Objective:** In this study, we investigated PPN activities during gait to better understand its functional role in locomotion. Specifically, we investigated whether PPN activity is rhythmically modulated during locomotion.

**Methods:** PPN local field potential (LFP) activities were recorded from PD or MSA patients suffering from gait difficulties during stepping in place or free walking. Simultaneous measurements from force plates or accelerometers were used to determine the phase within each gait cycle at each time point.

**Results:** Our results showed that activities in the alpha and beta frequency bands in the PPN LFPs were rhythmically modulated by the gait phase within gait cycles, with a higher modulation index when the stepping rhythm was more regular. Meanwhile, the PPN-cortical coherence was most prominent in the alpha band. Both gait-phase related modulation in the alpha/beta power and the PPN-cortical coherence in the alpha frequency band were spatially specific to the PPN and did not extend to surrounding regions.

**Conclusions:** These results raise the possibility that alternating PPN stimulation in tandem with the gait rhythm may be more beneficial for gait control than continuous stimulation, although this remains to be established in future studies.

## Introduction

The pedunculopontine nucleus (PPN) is a reticular collection of neurons at the junction of midbrain and pons.^1,2^ Its main mass is part of the mesencephalic locomotor region (MLR) in the upper brainstem, and it is thought to play an important role in modulating posture, locomotion, and arousal.^3,4,5^ Electrical stimulation of the MLR that directly projects to the locomotor central pattern generators in the spinal cord can generate walking, trotting, and galloping in a decerebrated cat^6^ and transition between swimming-like and walking-like movements in the salamander.^7^ Deep brain stimulation (DBS) of the PPN has been proposed as an emerging treatment for patients with Parkinson’s disease (PD) with gait problems,^8,9^ and for patients with multiple system atrophy (MSA) suffering from gait-related atypical parkinsonian syndromes.^10–12^ However, the therapeutic efficacy of PPN DBS and the extent to which it can improve quality of life seem too variable to draw firm conclusions, rendering it still an experimental therapy.^13^ This could be due to the still unclear complex structure-function relationships between these structures and locomotion.^14,15^

Conventional DBS settings deliver electrical pulses continually at a fixed frequency to the target brain area. Considering the rhythmic structure of movements during gait, conventional continuous DBS may be suboptimal for improving gait control.^16^ Understanding how neural activity in the PPN is modulated during gait could be a crucial step in improving the therapeutic effects of PPN DBS. Increased alpha power in the PPN has been observed in human patients during locomotion including free walking or imaginary gait,^3,17,18^ and increased alpha power in the PPN during walking correlated with gait performance.^17^ Similarly, increased low frequency oscillations in the PPN in the 1-8 Hz range have also been reported in a more recent study.^19^ However, the latter study reported a negative relationship between increased power in this low frequency band and movement speed especially when patients were off medication, which conflicts with some of the previous literature.

The previous literature so far has focused on the changes in the average power of the oscillatory activities in different frequency bands during walking compared with rest or standing still. Whether and how the activities in PPN LFPs are modulated within a gait cycle during walking or stepping are still unknown. This study aims to address this question as this may lead to new patterns of PPN stimulation that are more effective in ameliorating gait difficulties and facilitate rhythmic walking. To this end, we recorded PPN LFPs from patients undergoing DBS surgery targeting PPN for gait difficulties during stepping or free walking with simultaneous measurements from force plates or an accelerometer attached to the trunk to determine the phase within each gait cycle of each time point. Our results show that PPN activities in the alpha and beta frequency bands are rhythmically modulated by the phase of gait. The modulation is higher during more regular stepping, and the modulation is spatially limited to the PPN area.

## Methods

### Ethics

The present study was approved by the local ethics committees of the University of Oxford Hospitals or University College London Hospitals NHS foundation trust. All patients provided informed written consent before the experiment.

### Participants

Data were recorded from seven PD patients and four MSA patients who underwent bilateral (n = 9) or unilateral (n = 2) implantation of DBS electrodes targeting the PPN. The DBS electrodes were implanted and temporarily externalized (3-7 days) prior to a second surgery to connect the leads to a pulse generator in all patients. All experiments were conducted after overnight withdrawal of all dopaminergic medication. In total, data recorded from 20 PPN were analysed in this study (the data pertaining to average power changes during walking from all the seven PD patients (12 hemispheres) have been previously reported).^17^ The clinical details of the patients are summarised in Table I.

### Experimental protocol

A rest recording during which patients sat comfortably and relaxed with eyes open for 2-3 min was completed for each patient. Each PD patient completed another rest recording while standing for 2-3 min followed by a gait recording, where patients walked at their preferred speed along an unobstructed path (∼ 10 m) 10-30 times, depending on speed and fatigue. Each MSA patient completed a stepping in place recording during which they stepped on two pressure sensor plates (Biometrics Ltd, US) guided by a walking cartoon man displayed on a laptop while sitting comfortably in a chair, with a metronome sound provided at the time of each heel strike of the cartoon man, similar to the paradigm used in previous studies.^20,21^ Two MSA (MSA01, MSA02) patients with less severe gait problems were also recorded while stepping on the spot while standing. The participants were asked to follow the stepping rhythm of the walking cartoon man as precisely as possible, with one complete cycle (i.e., 1 right step and 1 left step) lasting 2 s.

### Recordings

For directional DBS leads, all segmented contacts of level 2 or 3 were physically joined together to make one monopolar ring contact. PPN LFPs and EEGs covering frontal and motor areas were recorded in monopolar configuration for all patients (no EEGs for PD07). In six PD patients (PD01-PD06), triaxial accelerometers were taped over the upper thoracic spinous processes to record their trunk accelerations. In all MSA patients, stepping force from left and right feet were recorded separately using two pressure sensor plates (Biometrics Ltd, US). All these signals were simultaneously recorded using a TMSi Porti amplifier (TMS International, Netherlands) with a sampling rate of 2048 Hz and common reference rejection.

### Data analysis and statistics

#### Pre-processing

The recorded data were first visually explored using Spike2 (v7.02a, Cambridge Electronic Design, UK) and those gait blocks/channels with obvious movement-related artefacts were rejected. The data were analysed offline with Matlab (v2020a, MathWorks, US). We first constructed bipolar LFPs using each pair of two spatially adjacent electrode contacts (e.g., 0-1, 1-2, and 2-3, where contact 0 was the most caudal) and quantified bipolar EEGs between “Fz” and “Cz”. Then, all bipolar LFPs and EEGs were band-stop filtered at 48-52 Hz, and band-pass filtered at 0.5-250 Hz. The recorded stepping force was band-pass filtered at 0.5-5 Hz.

#### Power spectral density and PPN-cortical coherence

The filtered bipolar LFPs and EEGs were decomposed into the time-frequency domain by applying continuous complex Morlet wavelet transforms. The wavelet power of each frequency was further normalized by the sum of the whole frequency band between 1-95 Hz, resulting in power spectral densities (PSDs) in percentage. In addition, imaginary coherence (IC) between each bipolar LFP and EEG (“CzFz”) was calculated for different movement states.^22,23^

#### Gait phase determination

To investigate the gait phase-related neural oscillations in the PPN, we determined the gait phase for each time point using the recorded force signals or accelerometer measurements. Specifically, when stepping force measurements were available, we first found all zero-crossing points from the band-pass filtered force data. Then, the zero-crossing points with the force decreasing, which corresponds to the time point when the contralateral foot started to lift up, were assigned as phase π or –π; the zero-crossing points with the force increasing, which corresponds to the time point of foot touching onto the force plate and starting to carry weight, were assigned as phase 0. The phases of all other time points were determined through linear interpolation between −π and 0 or between 0 and π, as shown in Fig. 1A. When stepping force measurements were unavailable, we used accelerometer measurements to manually identify each individual gait cycle. However, we only managed to identify the gait cycle in two patients (PD01 and PD02) because there was no clearly distinguishable rhythmic pattern in the accelerometer measurements during walking in the other Parkinson’s patients.

**Figure 1.**
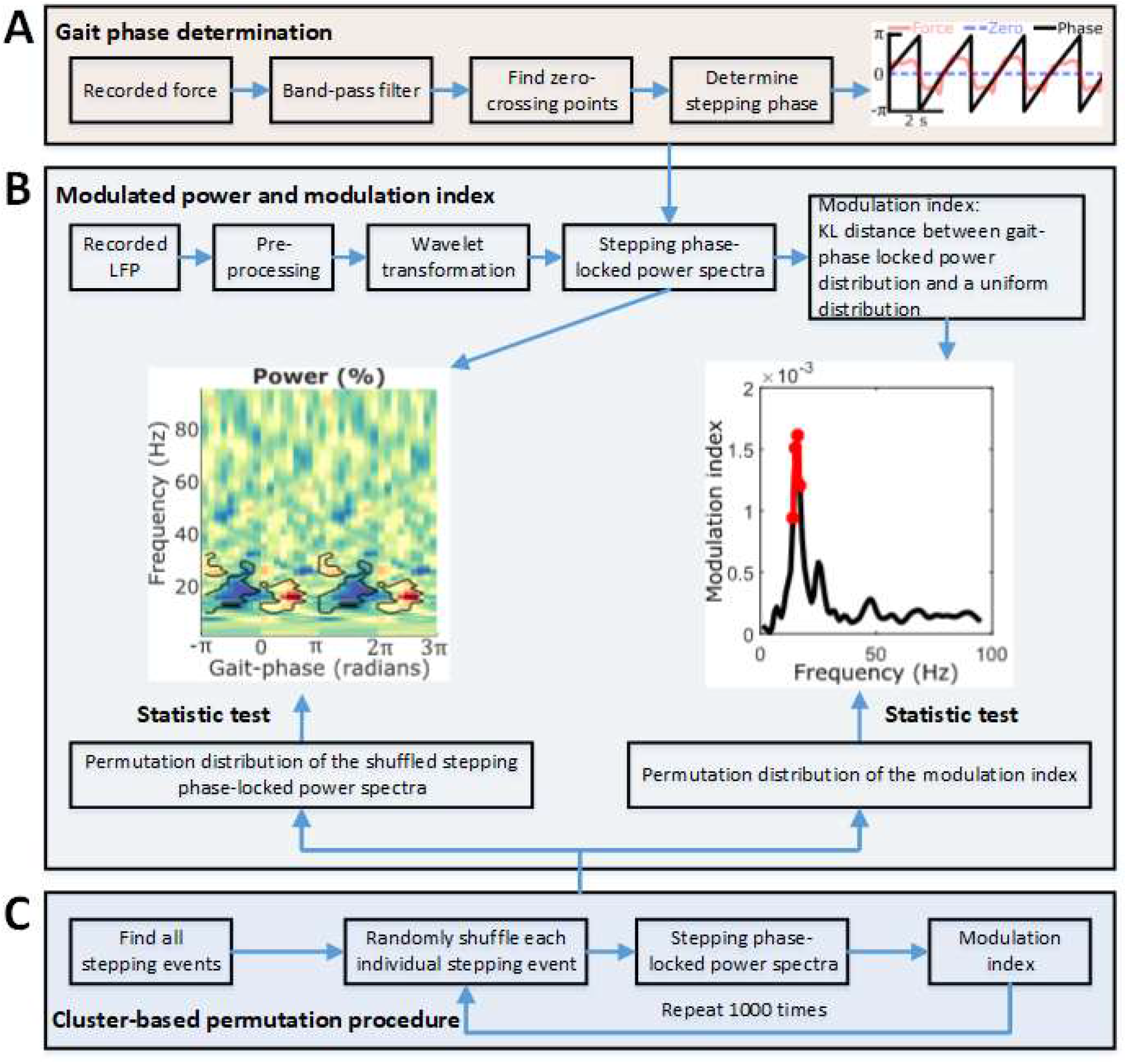
Flow chart for the data analysis procedure.

#### Power modulated by gait phase and the modulation index

Based on the wavelet power and the identified stepping/walking phase, we estimated the LFP activity-gait phase modulogram for each individual bipolar LFPs for the six patients (4 MSA and 2 PD). Specifically, 18 non-overlapping bins were defined from −π to π for each gait cycle. Then, the mean power within each phase bin was calculated for each frequency and normalized against the mean of all bins (Fig. 1B). A modulation index defined using the Kullback-Leibler (KL) distance between the gait-phase locked power at each frequency and a uniform distribution was also quantified for each frequency.^24,25^

#### Clustering of underlying states in PPN LFPs using Hidden Markov Model and K-medoids

We also investigated underlying states in the PPN LFPs defined by different dynamics in the LFP time series and how the state occupancy changes with gait phase using the HMM-MAR for each patient separately.^26^ In the HMM-MAR modelling, the maximum number of states was set to 30 and each time point would be assigned to a specific state. Then we quantified the occurrence rate of each state at different gait phase bins, and clustered these histograms into three main clusters using K-medoids to see how the occupancy of different states changes with the gait phase.^27^

#### Electrode trajectories reconstruction using Lead-DBS

To investigate the spatial distribution of the gait-phase related modulation, the electrode trajectories and location of different contacts were reconstructed using Lead-DBS based on preoperative MRI and postoperative CT scans for the five patients with identified gait phase (the scans for PD01 were not available),^28^ and the modulation index and PPN-cortical coherence during rest and gait for each individual bipolar contact were mapped into the MNI 152 NLIN 2009b space (Montreal Neurological Institute) using the Harvard AAN atlas.^29^

### Statistical analysis

A non-parametric cluster-based permutation procedure was applied to identify significant gait-phase related power modulation in the time-frequency plots while controlling for multiple-comparisons.^30^ To compare the group averages between two conditions, the condition labels of the original samples were randomly permutated 1000 times such that for each hemisphere the order of subtraction can change. To test the significance of points in the modulogram and the modulation index for individual bipolar contact recordings, we randomly shuffled the phase for each individual gait cycle and calculated the modulated power and modulation index. This was repeated 1000 times to obtain a permutation null-hypothesis distribution with 1000 samples (Fig. 1C). The permutation distribution mean and standard deviation were used to z-score the original unpermuted data and each permutation sample and get a *p*-value for each data point. Then, suprathreshold-clusters were obtained for the original unpermuted data by setting a pre-cluster threshold (*p*<0.05). If the absolute sum of the z-scores within the original suprathreshold-clusters exceeded the 95th percentile of the 1000 largest absolute sums of z-scores from the permutation distribution, it was considered statistically significant.

## Results

### PPN activity during rest and stepping

During standing or stepping compared with resting while sitting, the PPN activity and PPN- cortical coherence were modulated in the alpha and beta bands (Supplementary Fig. 1). Both in PPN LFPs and cortical EEGs, the power in the beta band was significantly reduced while standing compared with sitting, with peaks in the alpha frequency band observed in both conditions. The imaginary coherence between PPN LFPs and cortical EEGs was significantly reduced in the beta band (23-25 Hz, *t* = 6.1404, *p* = 0.014) and increased in the gamma band (57-65 Hz, *t* = −8.7731, *p* = 0.002) during standing. The results from MSA patients were similar, but the significant bands were narrower and the difference in gamma coherence was not significant.

### Modulation of PPN LFP activities within gait cycles

The LFP activity-gait phase modulogram and the modulation index were quantified for each of the individual bipolar PPN LFPs based on the stepping/walking phase of the contralateral foot. During stepping while sitting (Fig. 2A), stepping while standing (Fig. 2B), or free walking (Fig. 2C), significant modulation was observed in all hemispheres from all patients, with the strongest modulation in the alpha and beta bands. Note that PD01 received only a unilaterally implanted electrode and two bipolar channels for PD02 (L23 and R12) were excluded because of movement artefact in the LFPs during walking.

**Figure 2.**
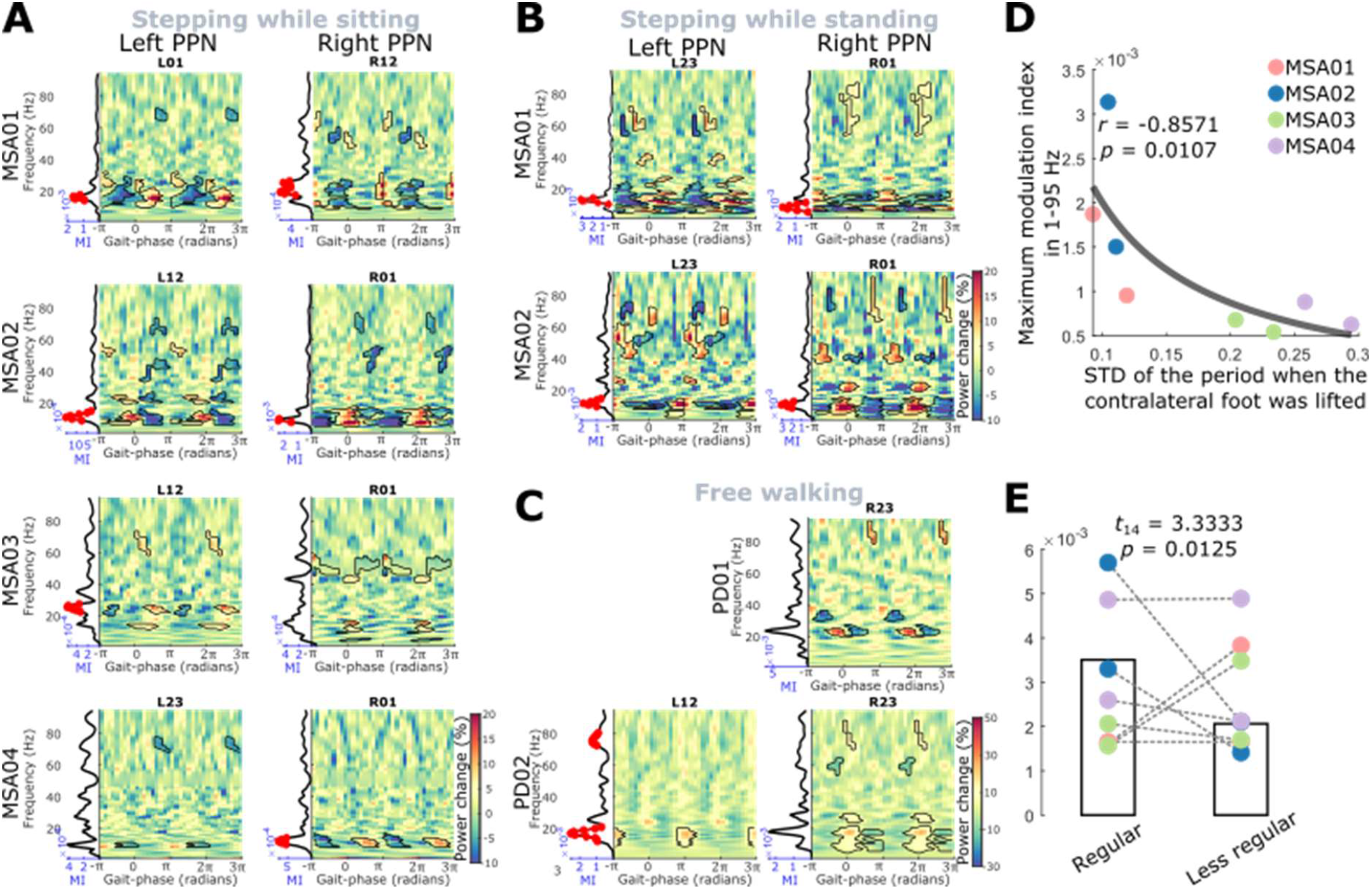
Power modulation during gait for the bipolar channel showing strongest modulation effect in each tested PPN. **(A)-(C)** Similar modulation was observed during stepping while sitting (A), stepping while standing (B), and free walking (C). In each subplot, modulogram (one bipolar channel from each hemisphere) showing the power in PPN LFPs modulated by the gait phase, with zero indicating the heel strike and −π (π) indicating the heal lift of the contralateral foot. The line plot to the left of the modulogram shows the corresponding modulation index. The black contours and red dots indicate significant modulation based on a cluster-based permutation procedure. **(D)** Relationship between the max modulation index and the variability of the stepping while sitting. Y axis indicates the maximum modulation index in a wide frequency band (1-95 Hz) and X axis indicates the standard deviation (STD) of the period when the contralateral foot was lifting in the air. Different colours indicate the point for different patients. The grey curve is a 1/x fitting of all data points. **(E)** The maximum modulation index during more regular stepping was significantly bigger than during less regular stepping.

When averaging the time-frequency plots and the modulation indices across all LFP channels for all recorded hemispheres, higher modulation was observed in the alpha and beta bands with one peak within one gait cycle in the PPN (Supplementary Fig. 2). Gait-phase related modulation was also observed in cortical EEG (CzFz), with peak modulation frequency in the high beta range and two peaks in the beta band observed within one gait cycle in the cortical signals.

### More regular stepping tends to be associated with stronger gait-phase related modulation in PPN LFPs

To investigate the impact of stepping variability on the observed gait-phase related modulation in PPN LFPs, we first quantified the standard deviation (std) of the duration when the contralateral foot phase, which was determined on a cycle-by-cycle basis, was between 0 and π across all stepping cycles for all patients with MSA and correlated them with the maximum modulation index in a wide frequency band (1-95 Hz). As shown in Fig. 2D, the two patients with stronger modulation effect (MSA01 and MSA02) had a smaller stepping variability as well as smaller gait impairment scores indicating less severe motor impairment based on the unified MSA rating scores (UMSARS-II) (Table 1). Across the eight hemispheres we found a negative correlation between the stepping variability and the maximum modulation index (*r* = −0.8571, *p* = 0.0107, Spearman’s correlation). Then, we split all stepping cycles of each leg into a more regular group and a less regular group with 25% cycles in each group for each hemisphere and compared the maximum modulation index between these two conditions. As shown in Fig. 2E, the maximum modulation index was significantly higher during more regular stepping (*t*_14_ = 3.333, *p* = 0.0125, paired *t*-test). These results suggest that more regular stepping tended to induce stronger modulation in PPN.

**Table 1:** Patients details. All patients were operated in Oxford except PD01 (London). For all motor scales, higher scores indicate worse function. The MSA patients had been attributed to Parkinson’s disease before clinically diagnosed as MSA. G = gender; yr = year; DD = disease duration; DBS = deep brain stimulation; UPDRS III = the motor subsection (part III) of Unified Parkinson’s Disease Rating Scale; IT27/30 = items 27–30 of UPDRS assessing posture, gait and balance; UMSARS II = the motor examination scale (part II) of unified MSA rating scale; Abb = Abbott infinity 1.5mm spaced leads (1-4), Abbott; Met^1^ = model 3387 1.5 mm spaced leads, Medtronic; Met^2^ = model 3389 0.5 mm spaced leads, Medtronic; SD = standard deviation. NA: Not available.

|  | G, age | DD<br>(yr) | DBS<br>lead | UPDRS III<br>OFF/ON<br>medication<br>(score/108) | IT27/30<br>OFF/ON<br>medication<br>(score/16) | UMS<br>ARS<br>II | L-dopa<br>dose<br>equivalent<br>(mg/day) | Other Pre-Op Medication | Data & Task |
| --- | --- | --- | --- | --- | --- | --- | --- | --- | --- |
| PD01 | M, 71 | 20 | Met<sup>1</sup> | 37/19 | 10/5 | NA | 1450 | NA | Rest sitting; Rest standing; Free walking |
| PD02 | M, 55 | 25 | Met<sup>2</sup> | 33/22 | 6/5 | NA | 300 | NA | Rest sitting; Rest standing; Free walking |
| PD03 | M, 68 | 9 | Met<sup>2</sup> | 40/26 | 11/8 | NA | 1650 | NA | Rest sitting; Rest standing |
| PD04 | M, 55 | 14 | Met<sup>2</sup> | 35/24 | 7/6 | NA | 1600 | NA | Rest sitting; Rest standing |
| PD05 | M, 70 | 20 | Met<sup>2</sup> | 35/22 | 6/5 | NA | 900 | NA | Rest sitting; Rest standing |
| PD06 | M, 76 | 16 | Met<sup>2</sup> | 34/25 | 9/9 | NA | 600 | NA | Rest sitting; Rest standing |
| PD07 | M, 54 | 20 | Met<sup>2</sup> | 53/19 | 6/5 | NA | 800 | NA | Rest sitting; Rest standing |
| MSA01 | M, 74 | 2 | Abb | NA | NA | 16 | NA | Ropinirole 14mg once a day | Rest sitting; Step sitting; Step standing |
| MSA02 | M, 72 | 2 | Abb | NA | NA | 12 | NA | Co-beneldopa 50/12.5mg 2 capsules 3 times a day | Rest sitting; Step sitting; Step standing |
| MSA03 | M, 71 | 2 | Abb | NA | NA | 17 | NA | Co-carbeldopa 25mg/100mg tablets: Dose: 1 tablet 3 a day | Rest sitting; Step sitting |
| MSA04 | M, 54 | 1.5 | Abb | NA | NA | 27 | NA | Co-beneldopa 25mg/100mg tablets: Dose: 2 capsules 4 times a day. Rasagiline 1mg once a day. Amantadine 100mg twice a day. | Rest sitting; Step sitting |
| Mean | 65.45 | 11.95 | NA | 38.14/22.43 | 7.86/6.14 | 18 | 1042.86 | NA | NA |
| SD | 8.51 | 8.53 | NA | 6.42/2.56 | 1.96/1.55 | 5.52 | 488.75 | NA | NA |

### Occurrences of different LFP states identified using HMM were modulated by gait phases

The unsupervised machine learning algorithms (MAR-HMM and K-medoids clustering) identified different clusters of PPN LFP states characterised by the multiple autoregressive modelling of the time series of the LFPs,^26^ with highest occurrence at opposite points of the stepping/walking phases (Fig. 3 A+B and D+E). Specifically, states in cluster 1 were more likely to occur at phases −π and π (when the foot is lifted), and at phase 0 (at the time of the heel strike). The occurrence of cluster 2 states was most probable at −π/2 (when the foot was highest in the air) and π/2 (in the middle of the stance period). A similar pattern was detected in the PD patient (PD02). We did not include PD01 here because the number of recorded walking cycles (25) was too small to provide reliable state estimates.

**Figure 3.**
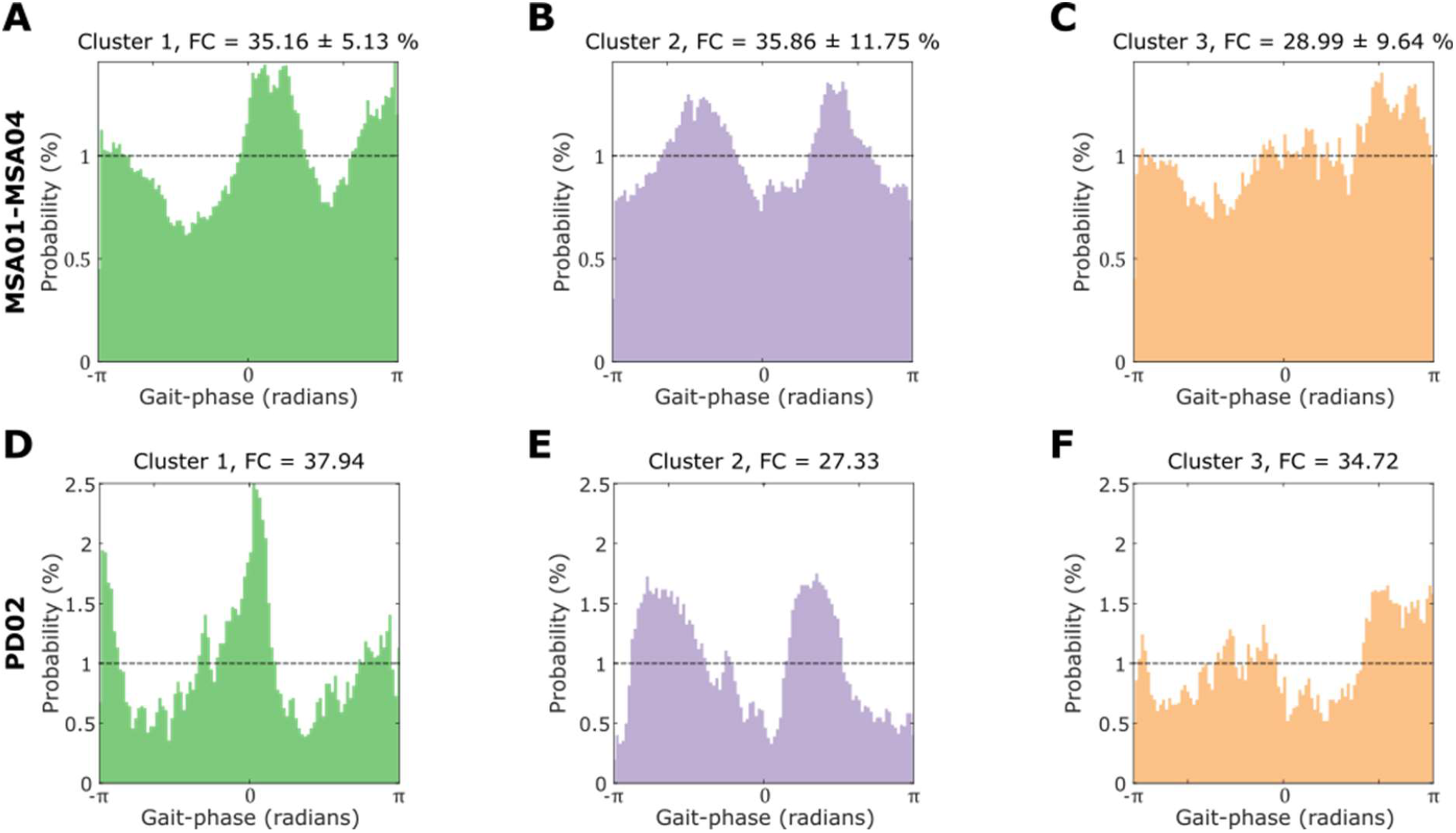
Clusters with different occurrence rate at different gait phases were identified using unsupervised machine learning algorithms (HMM and K-medoids clustering). **(A)** Cluster 1 showed states with higher probabilities at phase −π/π (when the foot was lifted) and at phase 0 (at the time of the heel strike) during stepping while sitting. **(B)** Cluster 2 showed higher probabilities at phase −π/2 (when the foot was highest in the air) and π/2 (in the middle of the stance period) during stepping while sitting. **(C)** Cluster 3 did not show specific preferred gait-phases during stepping while sitting. The results in (A)-(C) were averaged across all four MSA patients. (D)-(E) Very similar results were obtained for one PD patient during free walking. Please note that each histogram consisted of 100 phase bins. The black dash line indicates the equally probability (i.e., 1%) of each phase bin if there was no modulation effect.

### Alpha and beta gait-phase related modulation were clustered in the PPN

Electrode location reconstruction using the LeadDBS software confirmed that most electrodes were well placed in the PPN area with respect to the Harvard AAN atlas,^29^ as shown in Fig. 4A. The gait-related power modulation was mainly observed in alpha and beta frequency bands but not in theta and gamma frequency bands (Fig. 4B). There was a wider spatial spread of gait-phase related modulation in the beta band activities during stepping or walking; in comparison, the modulation in alpha band activities was more focused close to the PPN. In addition, the imaginary coherence between PPN LFPs and cortical EEGs during rest or gait mainly occurred in the alpha band, but not in beta, theta or gamma bands (Fig. 4C, D).

**Figure 4.**
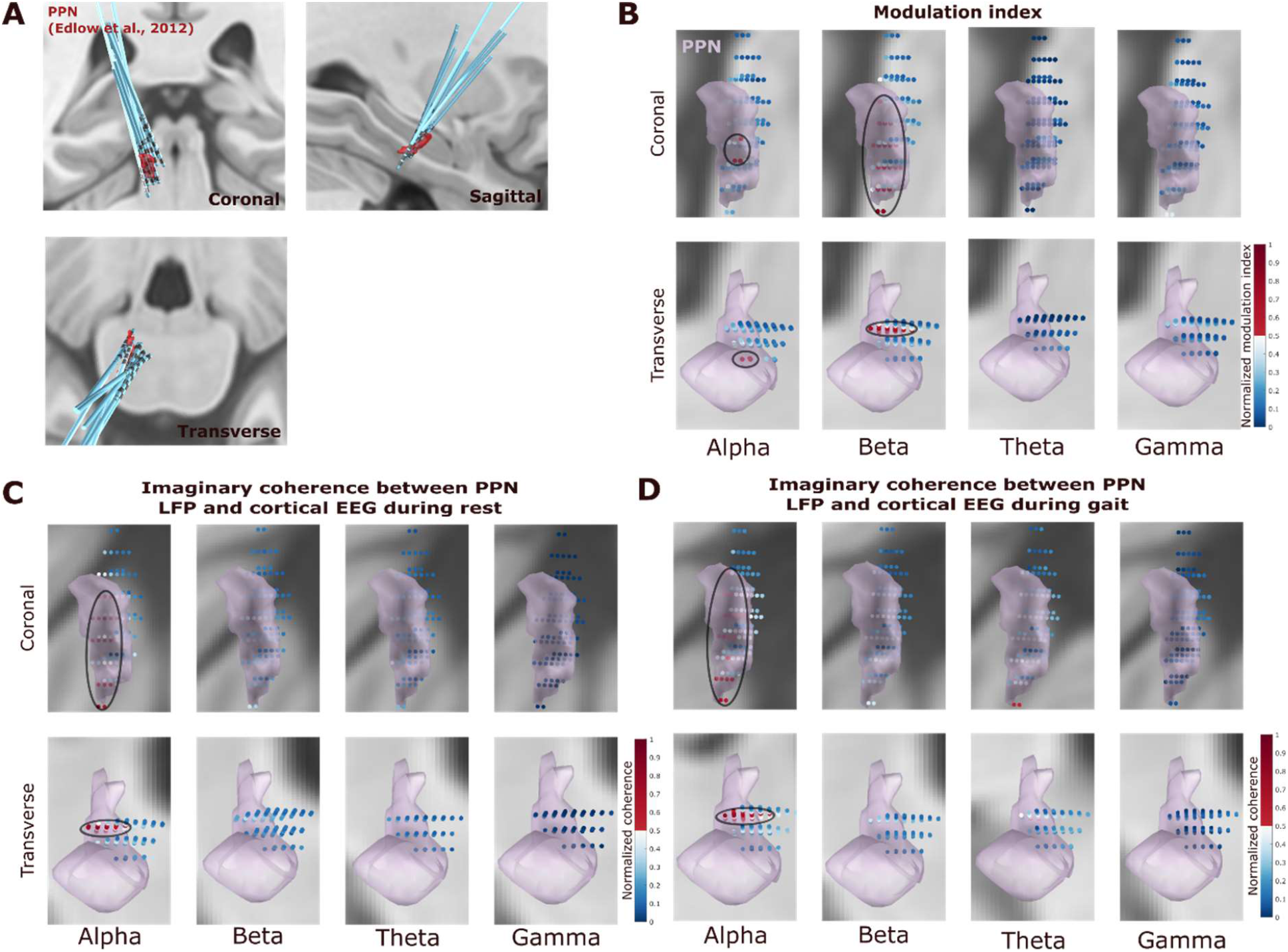
Electrode localization (A) and the mapping of the modulation index (B) as well as the PPN- cortical coherence (C) in standard MNI space. **(A)** 3D reconstruction in coronal (top left), sagittal (top right), and transverse (bottom left) views of all the recorded DBS leads using Lead-DBS software. **(B)** The bipolar contacts close to the PPN in the MNI space tend to show higher gait-phase related modulation index in alpha (8 - 12 Hz) and beta (13 - 30 Hz) frequency bands, and the modulation index in these two frequency bands were stronger compared with theta and gamma frequency bands. **(C)** The imaginary coherence between PPN LFP and cortical activities (measured at CzFz using EEG) averaged in the alpha frequency band was clustered close to PPN in the MNI space and was strongest compared with beta, theta, and gamma frequency bands. **(D)** The same holds for the imaginary coherence between PPN LFP and cortex EEG during gait. Black ovals indicate clusters of larger values. The two rows show the results in two different views.

## Discussion

We found that alpha and beta oscillations in the PPN LFP are modulated relative to the contralateral foot gait phase within gait cycles in patients with MSA and PD during stepping movements, no matter seated, or standing or during free walking. Gait phase-related modulation increased with more regular stepping movements, was specific to alpha and beta frequency bands, and clustered around the PPN area defined by the Harvard AAN atlas. These results are similar to those we have observed in STN.^20^ The gait-phase related modulation in the STN was strongest in the high beta band between 20-35 Hz,^20^ whereas gait-phase modulated PPN activity was specific to the alpha and low beta range. We are confident that the modulation reflects neural activity and not movement-related artefacts as artefacts are minimal during stepping while sitting^20^ and the modulation pattern was specific to limited frequency bands. These results are also in line with reports that alpha oscillations in the PPN are associated with gait in PD patients.^17,18,31,32^ However, previous studies hypothesized that increased alpha activity in the PPN is important for normal gait, and that low frequency PPN DBS supports or emulates this activity, consequently enhancing the allocation of attentional resources and improving gait performance.^2,13^ Here we provide novel evidence for gait-phase related modulation of the PPN activities in the alpha and beta bands for regular stepping. These modulations are more likely related to the movements rather than the allocation of attentional resources during gait, although the latter could modulate the gain of these cyclic modulations.

### Location

The boundaries of the PPN are still indistinct and there is inconsistency in the location of the PPN in different atlases.^2^ In this study, we utilized the PPN atlas defined in Lead DBS according to a study of Edlow et al. to reconstruct the trajectories of the DBS electrodes.^29^ With this atlas, we showed that the gait-phase related modulation was more focused in the ventral-rostral plane of the PPN in the alpha band and exhibited a wider spatial spread in the dorsal-caudal PPN plane in the beta band (Fig. 4B, Supplementary Fig. 3). It should be noted that the definition of the PPN borders in standard MNI space in other atlases maybe slightly different compared with the one we were using.^33–35^ In this study, we chose the atlas occupying the largest volume in the standard space. Nevertheless, since precise localization of PPN is difficult, the structures defined in all these atlases should be considered presumptive, with the atlas we chose being the most inclusive one.

### Potential implications in PPN DBS

Recently, an alternating DBS pattern, consisting of rhythmic intermittent reductions in stimulation intensity with a fixed offset between the right and left STN, was shown to significantly manipulate step timing.^16^ These results have raised the possibility that alternating stimulation in the STN may be a promising DBS pattern for gait control in PD. In this study, we showed that alpha and beta oscillations in the PPN were modulated relative to the gait phase of the contralateral foot in both patients with MSA and PD. The PPN has specific relevance to locomotion, as it is considered a key component of the MLR – an area where electrical stimulation in decerebrated animals can induce locomotor-like activity.^36^ It has widespread reciprocal connectivity with many structures including basal ganglia nuclei, thalamus, nuclei of the pontine and medullary reticular formations, deep cerebellar nuclei, and the spinal cord.^15^ Thus the PPN may be a more powerful target for alternating DBS than the STN in entraining rhythmic movements required for normal gait. Further studies as to how PPN DBS affects activities in the PPN and motor network, how stimulation effect changes with stimulation frequency,^13^ and how alternating PPN DBS might affect stepping performance are warranted.

### Limitations

The first limitation of this study other than the uncertainty in PPN spatial limits considered above is our small sample size. We analysed data from two PD patients and four MSA patients to quantify gait-phase related modulation in the PPN LFP. PPN DBS is still an emerging experimental treatment for patients with gait problems, and the number of patients operated on is small.^13^ The patients with MSA were involved in a recently registered clinical trial to investigate the clinical effect of PPN DBS on gait control. Due to the strict inclusion criteria of the trial and the impact of the pandemic, we were only able to record four patients in this cohort. However, the results were consistent within this small sample size and modulation was consistently observed across stepping movements, made while seated, standing or free walking. The second limitation is that this study is only correlational. It is impossible to distinguish whether the modulation we observed reflects motor output or sensory feedback about the movements. To test the causal role of rhythmic modulation of PPN activity and its potential clinical benefits, we need to test alternating PPN DBS patterns and evaluate their impact on gait control, similarly as previously done with alternating STN DBS.^16^

## Conclusion

This study provides new evidence that PPN activities in the alpha and beta bands are modulated by gait phase within gait cycles, with the modulation increased during more regular stepping movements. Our observations raise the possibility that modulating PPN DBS relative to the gait cycle mimicking the modulation pattern we observed during stepping could potentially facilitate the observed activity pattern and be more beneficial than continuous stimulation for facilitating the rhythmic movements required in normal gait.

## Acknowledgments

We thank the participating patients for making this study possible.

## Author Roles

1. Research project: A. Conception, B. Organization, C. Execution
2. Statistical Analysis: A. Design, B. Execution, C. Review and Critique
3. Manuscript: A. Writing of the first draft, B. Review and Critique

S.H.: 1C, 2B, 2C, 3A, 3B.

A.D.: 1C, 2C, 3B.

P.F.: 1A, 1B, 1C, 2A, 3B.

C.W.: 2C, 3B.

Y.H.: 1B, 1C, 2B, 2C.

S.M.: 2C, 3B

S. K.: 2C, 3C

T.Z.A.: 1A, 1B, 2A, 3B

A.L.G.: 1A, 1B, 2A, 2B, 3B

P.B.: 1A, 1B, 2A, 2B, 3B.

H.T.: 1A, 1B, 1C, 2A, 2B, 2C, 3B.

## Supplementary materials

**Supplementary figure 1.**
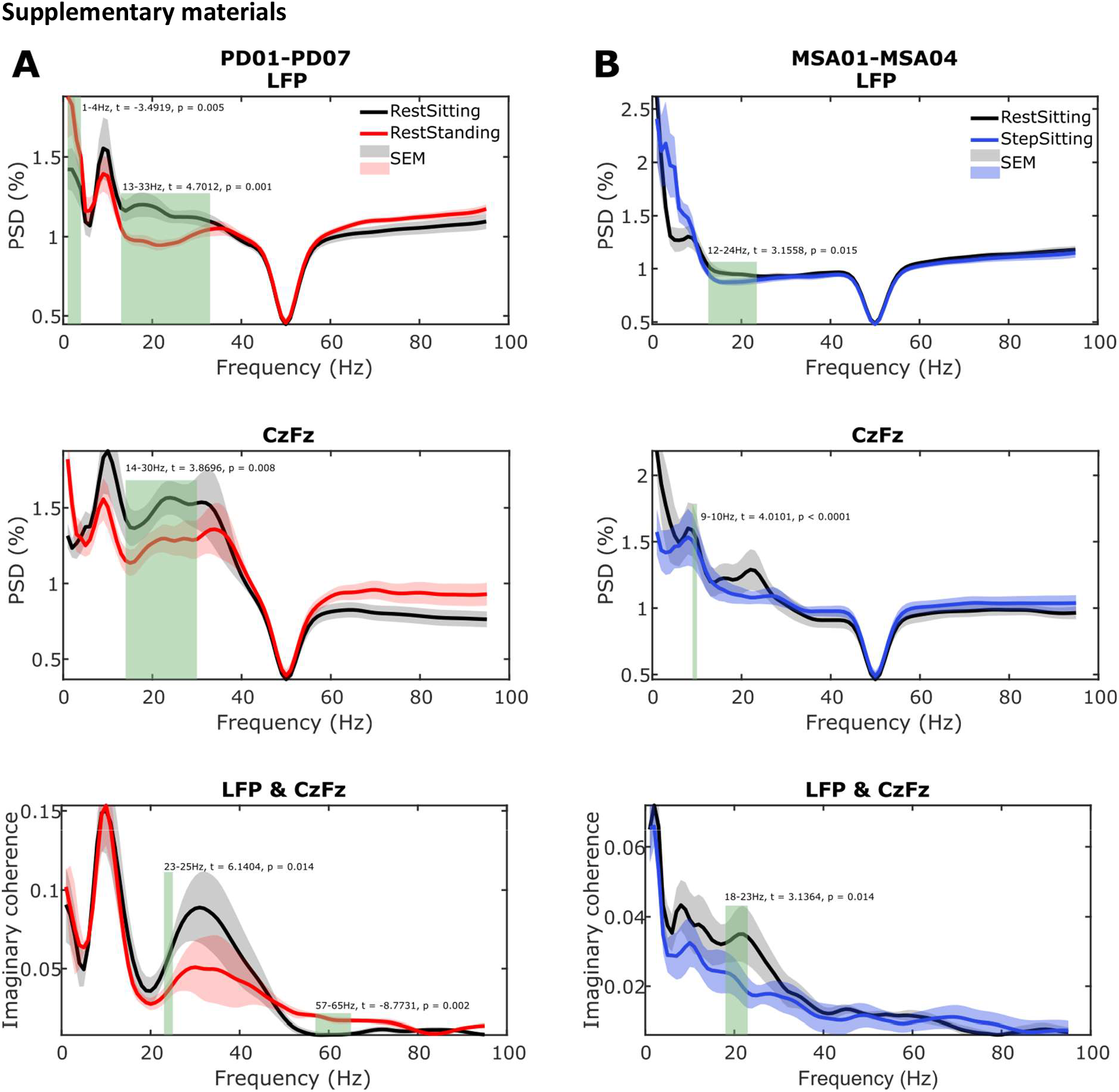
Power spectral density and coherence between LFP and EEG. **(A)** Power spectral density (PSD) of PPN activity (top), cortical EEG activity (middle), and imaginary coherence (IC, bottom) between PPN and cortical EEG during rest while sitting (black) and rest while standing (red) averaged across 7 PD patients (12 hemisphere). **(B)** PSD and IC during rest while sitting (black) and stepping while sitting (blue) averaged across 4 MSA patients (8 hemispheres). Green rectangle indicates significant difference based on a cluster-based permutation procedure.

**Supplementary figure 2.**
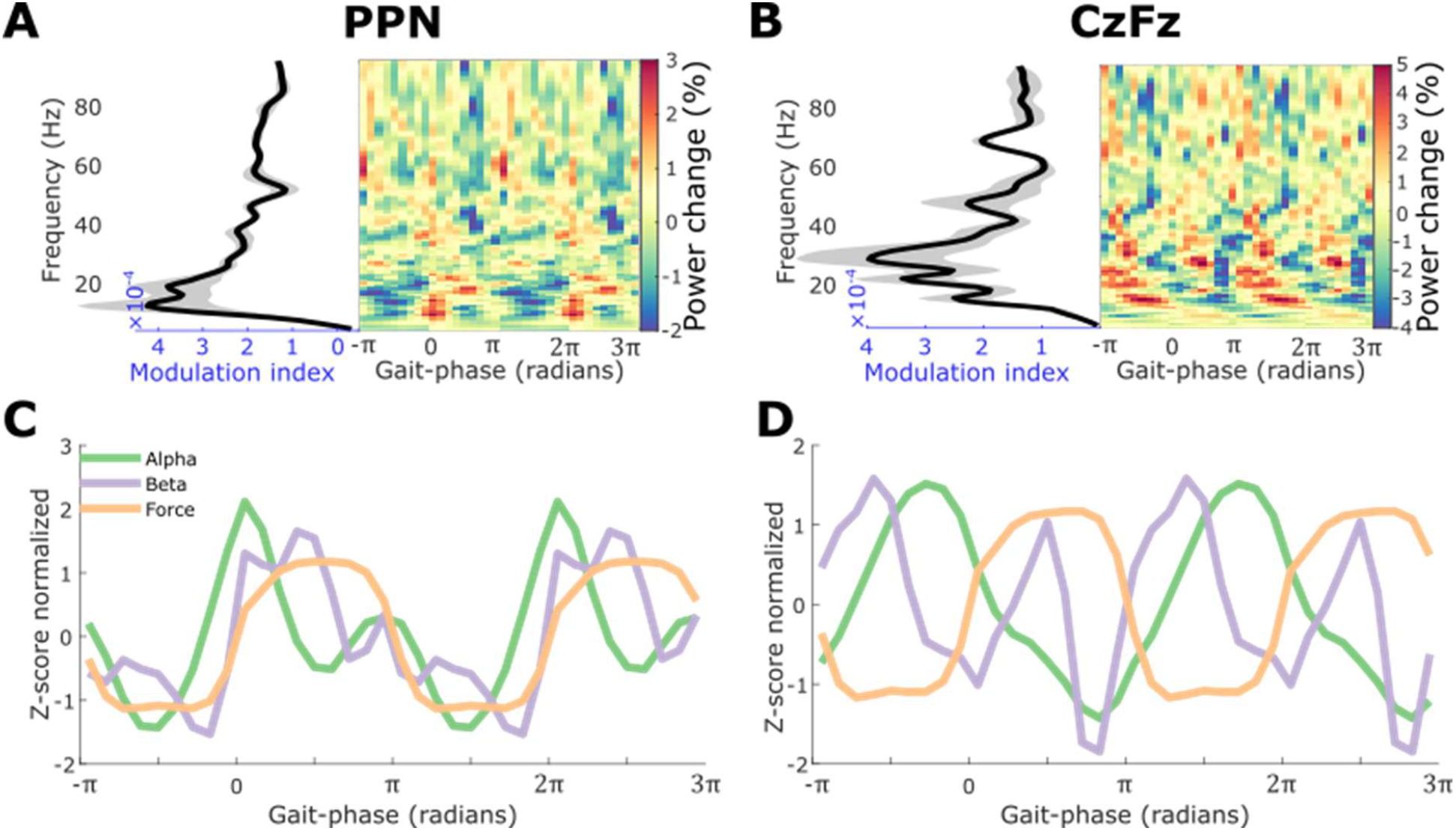
Averaged power modulation during stepping while sitting (n= 8 PPN of 4 MSA patients). **(A)-(B)** Power was modulated in both PPN LFP (A) and cortical EEG (B). **(C)-(D)** Averaged power in alpha (green) and beta (purple) frequency bands, and the averaged force (orange) relative to gait-phase in PPN (C) and cortical EEG (D).

**Supplementary figure 3.**
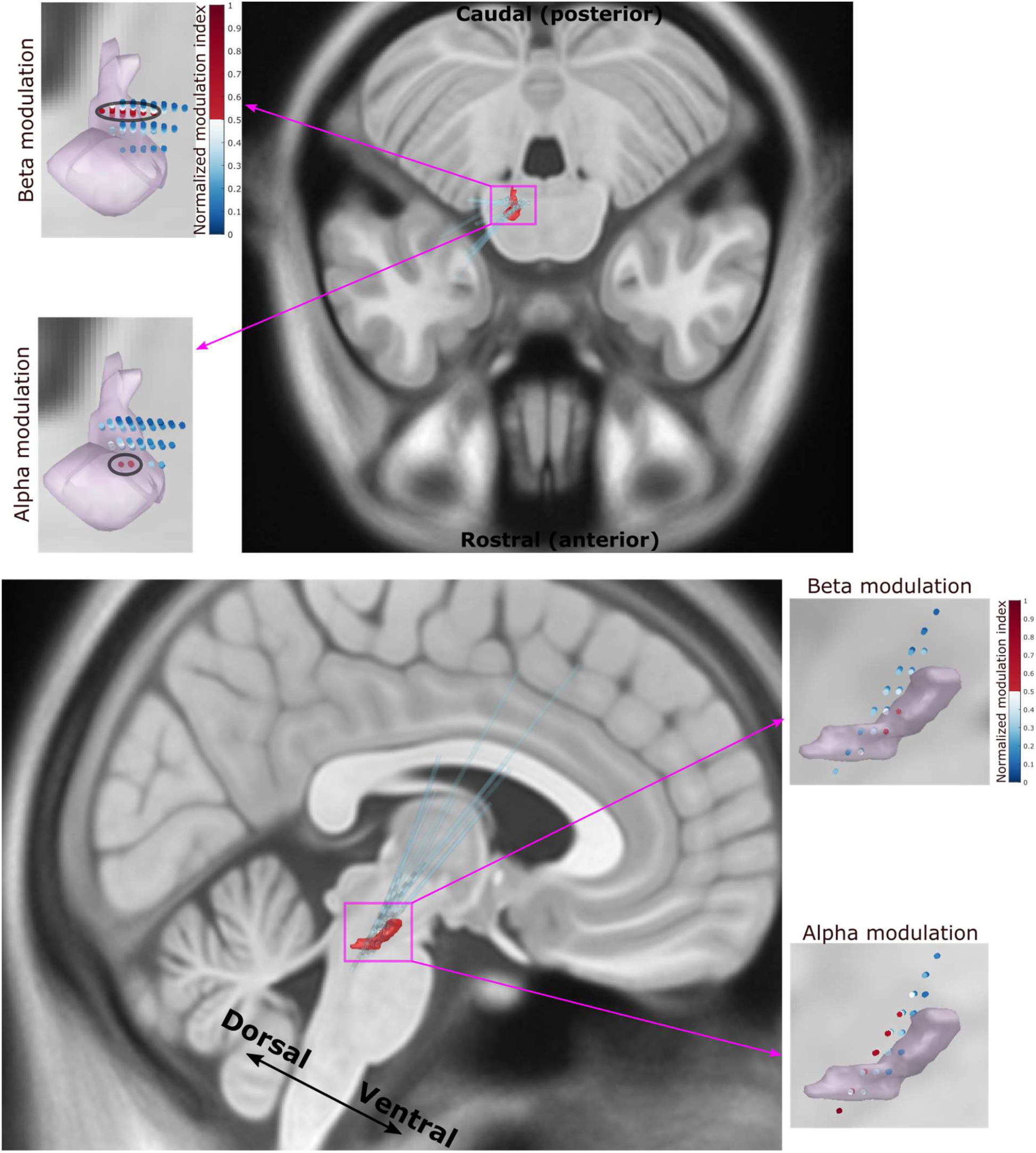
The strongest modulation effect in alpha band was more focused in the ventral-rostral portion of the PPN while the strongest modulation in beta band was more widely spatial spread in the dorsal-caudal PPN.

